# A double-edged sword: supramolecular complexes of triazavirine display multicenter binding effects which influence aggregate formation

**DOI:** 10.1101/305615

**Authors:** Yana A Zabrodskaya, Alexey V Shvetsov, Vladimir B Tsvetkov, Vladimir V Egorov

## Abstract

Triazavirine (TZV), a small molecule with a broad spectrum of antiviral activity, shows a tendency to induce the aggregation of melittin (MLT), a cationic membrane-disrupting peptide from the poison of the European bee. In MLT:TZV mixtures, high molecular weight precipitates form once a critical MLT:TZV ratio has been exceeded. Molecular dynamics simulation of MLT-TZV interactions, followed by molecular docking, led us to the hypothesis of multicenter binding of TZV supramolecular complexes to MLT.

## Introduction

MLT is a peptide from the poison of a European bee. Peptide oligomers are capable of forming pores in membranes of various cell types, in particular, red blood cells [1], [2]. In a number of studies, this effect on cells is compared with the effect of amyloid-like oligomers on cell membranes [3], [4] and, in general, membrane-destroying peptides. Therefore, studies regarding the modulation of membrane permeability with low toxicity substances can potentially aid in the development of drugs targeting diseases in which the destruction of membranes by exogenous or endogenous peptide oligomers play a role.

In the course of studies on TZV activity using model peptides by our colleague, Dr. Olga Alexandrovna Mirgorodskaya, an observation was made that the drug stimulates the precipitation of the peptide from solution. Because this observed TZV effect was contrary to what we previously documented when studying its effects on a peptide comprising an ionic self-complementary motif [5] [6] we decided to analyze the effect in detail. The formation of strong hydrogen bonds between arginine and TZV (mentioned in [5]) could explain aggregation of the generally hydrophobic peptide by a mechanism in which there is shielding of the electrostatic repulsion between arginines. The ability of MLT to oligomerize under the influence of TZV and the effect of the resulting oligomers on membranes was the subject of the study.

## Materials and Methods

### Materials

Human red blood cells (2% in PBS, phosphate buffered saline) were kindly provided by the influenza vaccine laboratory at the Research Institute of Influenza, St. Petersburg, Russia. Triazavirine (TZV; 2-methylthio-6-nitro-1,2,4-triazolo[5,1-c]- 1,2,4-triazine-7(4I)-one) was synthesized at the Ural Federal University in honor of B.N. Yeltsin, Ekaterinburg, Russia. The MLT Peptide and all other reagents were purchased from Sigma Aldrich.

### Atomic force microscopy

MLT at a concentration of 1 mM was mixed with TZV at different ratios, namely: 1:1; 1:5; 1:15; and 1:30. After 1h or 24h incubation, the samples were 10-fold diluted and put on freshly cleaved mica. After 1 min. incubation at room temperature, the mica surface was washed three times with water to remove salt. The sample topography measurements were performed in semi-contact mode on an atomic force microscope (NT-MDT–Smena B) using the NSG03 probe (NT-MDT, Russia). The images were analyzed using Gwyddion software [7].

### Congo Red assay

Congo red dye (Sigma Aldrich, USA) at a concentration of 50 μM in PBS was mixed with the 10 μM peptide. As a control, a similar sample was used in which a buffer was added instead of the peptide. Absorption spectra were recorded on an Avantes Ava Spec 2048 instrument (Avantes, The Netherlands).

### Red blood bell assay

MLT (10 μl, 0.04 mM in H_2_O) was pre-mixed with 10 μl TZV at different molar ratios: 1:1; 1:5; 1:15; 1:30; and 1:60. As a control, 10 μl of H_2_O was used instead TZV. After 15 min. of incubation at room temperature, the mixtures were centrifuged on a MicroSpin (Biosan). Human red blood cells (2% in PBS), 30 μl, were mixed with 10 μl MLT-TZV solution supernatant. These red blood cell and MLT concentrations were chosen because the measured absorbance at 415 nm indicated that the degree of erythrocyte lysis was linearly correlated with MLT concentration. The absorbance was measured on a NanoDrop ND-1000 spectrophotometer.

### Molecular dynamics simulation

The initial MLT structure was taken from a reference resource [8]. After that, 16 MLT peptide chains were inserted into a 1000 nm^3^ (10x10x10 nm) cell in random orientations using gmx insert-mol. In models which included TZV in addition to 16 MLT molecules, 320 TZV molecules were inserted with random orientations using gmx insert-mol. TZV modelling was accomplished as described previously [6].

All MD simulations were performed in the GROMACS [9], [10] software package using an amber99sb-ildn [11] force field for peptides, and a tip3p [12] model for explicit water. The simulated systems were around 140k atoms in size including an explicit solvent shell and 50 mM NaCl to neutralize charges on the peptides (and TZV, if present in the model). Solvent shell thickness was at least 2.5 nm. The systems were equilibrated using a two-step protocol. During the first step, the system was equilibrated for 5 ns with all heavy, non-solvent atoms restrained to their initial positions using NPT ensemble. During the second step, the system was equilibrated without restraints for 10 ns (starting from the last frame of the previous step). After two equilibration stages, a 500 ns trajectory was simulated with a time step of 2 fs. A Neighbour search was performed every 50 steps. The PME algorithm was used for modeling electrostatic and Van der Waals interactions with a cutoff of 1.2 nm. Temperature coupling was done with the Nose-Hoover algorithm at 310K. Pressure coupling was done with the Parrinello-Rahman algorithm for 1 bar.

All further calculations (pairwise distances, average minimal distances, occupancy) were performed on the last 100 ns time interval.

### Molecular docking

The three dimensional models of TZV and MLT were built using a molecular graphics software package, Sybyl-X (Certara, USA). In order to build the 3D model for MLT, the position for the atomic coordinates were taken from PDB:2MLT [8]. Partial charges on the ligands’ atoms were calculated according to the following scheme. First, in order to find the most minimal conformation, scanning of the conformational space of the ligands was performed with the application of a molecular-mechanical approach and a Monte Carlo method using Molsoft ICM-Pro 3.8.6 [13], [14]. To calculate interatomic interactions, the mmff force field [15] was used at this stage. Further optimization of the conformation found at the first step was carried out using density functional theory (DFT) using the hybrid exchange-correlation functional B3LYP (Becke three-parameter (exchange); Lee, Yang and Parr (correlation)) [16]–[18] with basic sets 6–311G+ + (2d,2p). For the resulting conformation, in order to calculate the partial atomic charges CHELPG (Charges from Electrostatic Potentials using a Grid based method)) [19], based on fitting of molecular electrostatic potentials (MEP), was used. All quantum mechanics simulations were carried out using the Gaussian 09 program [20]. To define the most probable binding site of the ligands on the target surface, the flexible ligand docking procedure was performed in ICM-Pro 3.8.6 software. Before starting a docking procedure, the structures of targets and ligands were converted into an ICM object. According to the ICM method, the molecular models were described using internal coordinates as variables. The parameters needed for interatomic energy calculation and the partial charges for the atoms of targets were taken from the ECEPP/3 [21]. The biased probability Monte Carlo (BPMC) minimization procedure [22] was used for global energy optimization. A description of the binding energy scoring used in the docking and BPMC procedures is reported by R. Abagyan and M. Totrov [23]. Finally, the conformational stack obtained from the docking procedure was sorted by scoring function, and 100 best complexes were taken and again minimized using a conjugate gradient method with gradient tolerance criterion equal to 0.05 kcal·mol^−1^·Å^−1^ and the mmff force field. Next, the binding energy of each complex was estimated using the formula:

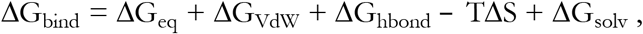

representing the sum of the free energy of complexation in a vacuum and ΔG_solv_, which is the difference between the solvation energy of the complex and the sum of solvation energy of the unbound target and the ligand. Energy complex formation in vacuum was evaluated as the sum of: the change in electrostatic ΔG_eq_; Van der Waals ΔG_VdW_ components; the energy of hydrogen bonds formation ΔG_hbond_; and TΔS (the change in entropy due to the loss of the conformational mobility of the target and the ligand). The solvation energy was evaluated by the Wesson–Eisenberg method [24]. The best complex was selected by using the results of estimation as it applies to the formula above. At the final stage, the selected complex was subjected to global energy optimization.

## Results and discussions

MLT was mixed with a solution of TZV in ratios from 1:1 to 1:60. At the same time, as observed by Mirgorodskaya, the formation of precipitates was documented. Analysis of precipitates by atomic force microscopy showed the presence of spherical aggregates in the mixture when TZV exceeded a certain limit (30 molecules of TZV per 1 molecule of MLT) (Figure 1, a-e). The size of the aggregates also depended on the MLT: TZV ratio and further incubation did not affect their sizes. It should be noted that samples with Congo Red dye, specific to amyloid-like fibrils [25], didn’t show the characteristic right-shift (data not shown).

**Figure 1.**
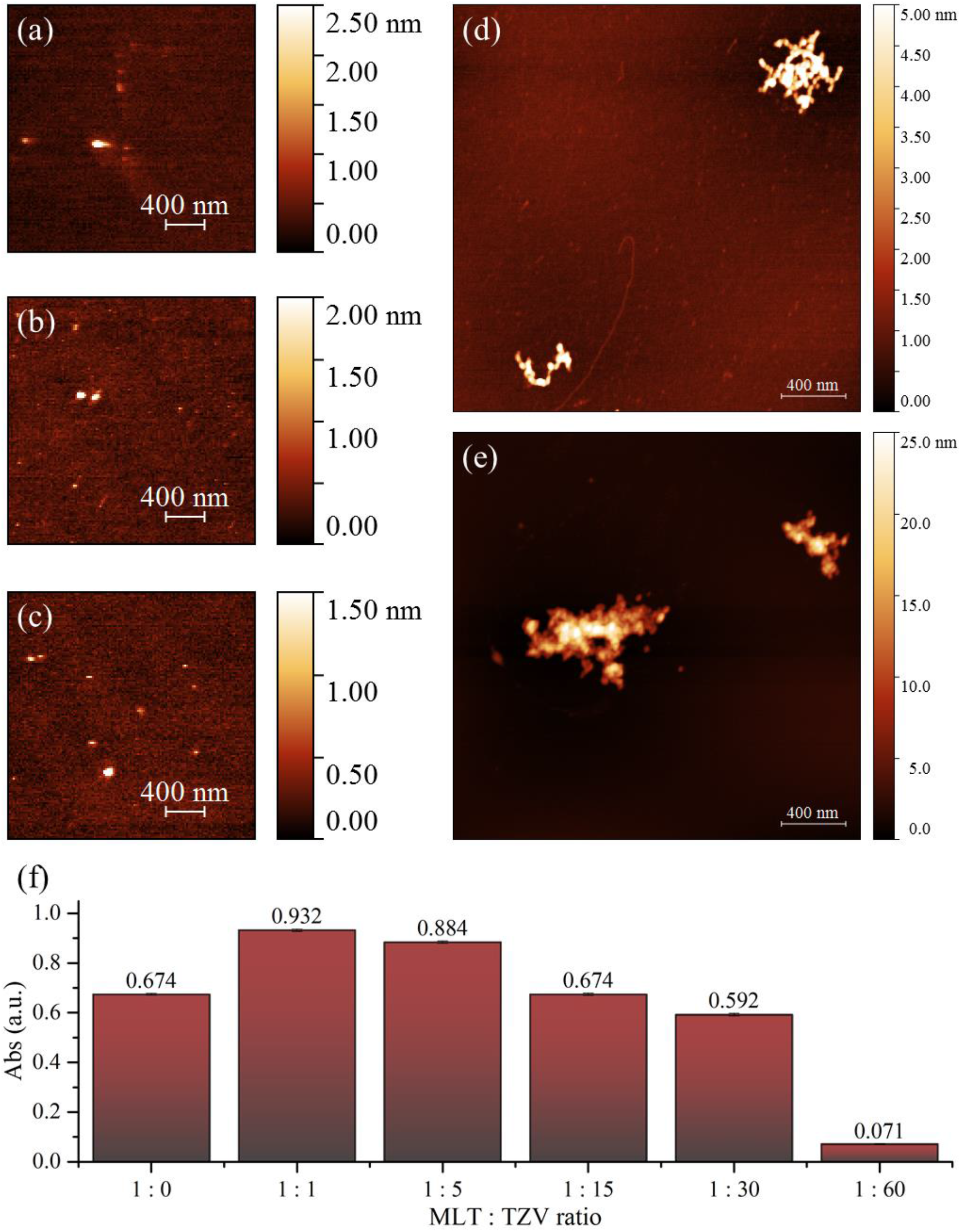
The surface topography of intact MLT **(a)** and its mixture with TZV in ratios 1:1 **(b)**, 1:5 **(c)**, 1:15 **(d)**, 1:30 **(e)**. The scale bar is 400 nm. On the right of (a)-(e) is the pseudocolor ruler indicating the particles’ height (nm). **(f)** The absorbance at 415 nm indicating erythrocyte hemolysis as it depends on the MLT: TZV ratio. Absorbance of pure erythrocytes without MLT is 0.059 ± 0.004 a.u.

The mixtures were centrifuged and the activity of MLT in supernatants was studied on erythrocytes isolated from healthy volunteers (Figure 1, f).

The membranolytic activity of MLT in the presence of TZV changed nearly instantaneously and only when TZV exceeded MLT by 30 or more fold (1:30, 1:60). Moreover, at ratios from 1:1 to 1:5, a small increase in the membranolytic activity of MLT was observed, while TZV itself did not act as a membrane protective agent at any of the concentrations tested. Since MLT can exist in solution in two forms, trimers and tetramers [26], [27], it is possible that TZV facilitates the MLT transition to a more active conformation. It should be noted that if the precipitates that formed are not centrifuged, the hemolysis effect persists.

The tendency of TZV to form supramolecular complexes, described by us for the first time in the citation [6], explains this behavior of oligomers.

We performed a molecular dynamics simulation of the system under study in an aqueous box with conditions similar to those in citation [6]. It was shown that free MLT exists as a monomer in solution, and it tends to form short, unstable chains (Figure 2a). Interestingly, MLT is clusterized under TZV influence (Figure 2b).

**Figure 2.**
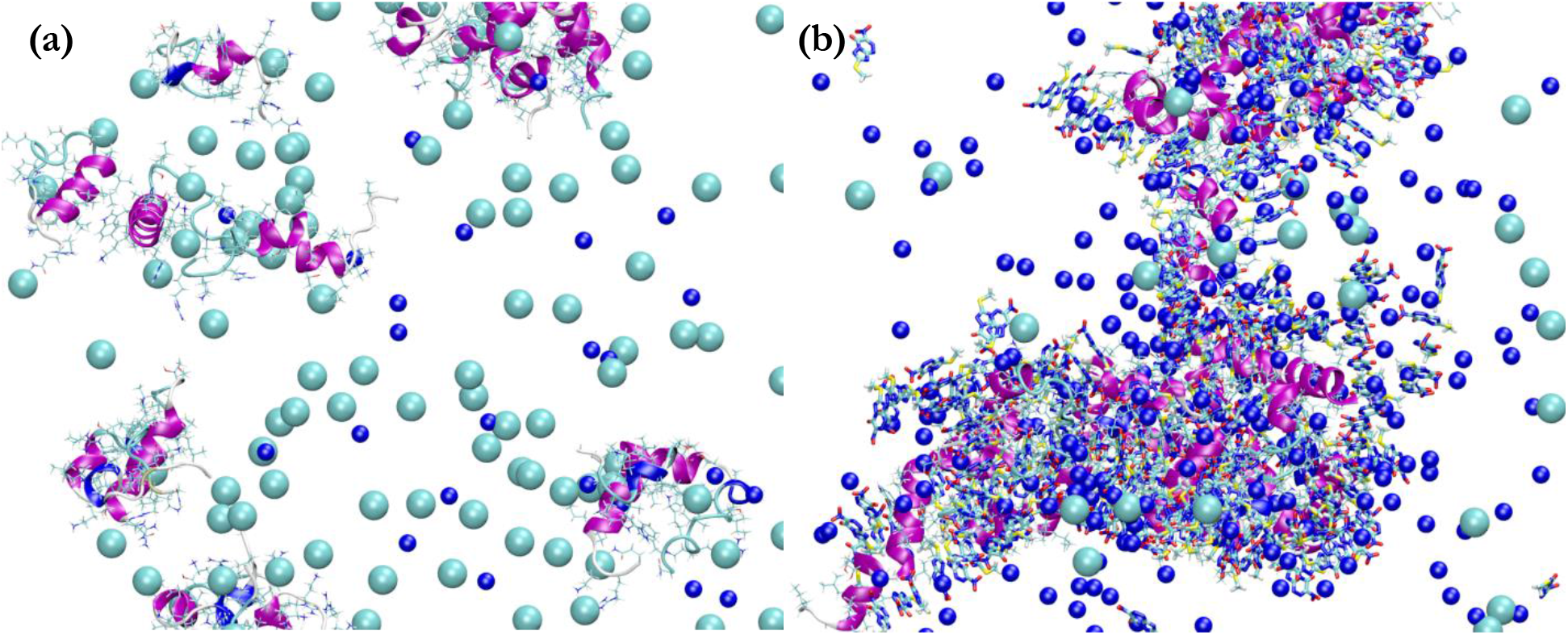
**(a)** Final state of a 500 ns molecular dynamics simulation of 16 MLT peptide molecules in solution and **(b)** 16 MLT peptide molecules in solution with 20-fold molar excess of TZV.

The simulation results showed that in the presence of self-organizing supramolecular chains of TZV, the pairwise distance between the centers of mass of the MLT molecules decreases (Figure 3a). It should be noted that some TZV molecules are strongly connected with MLT (Figure 3b), as seen by weak exchanges between bound TZV and free TZV in solution.

**Figure 3.**
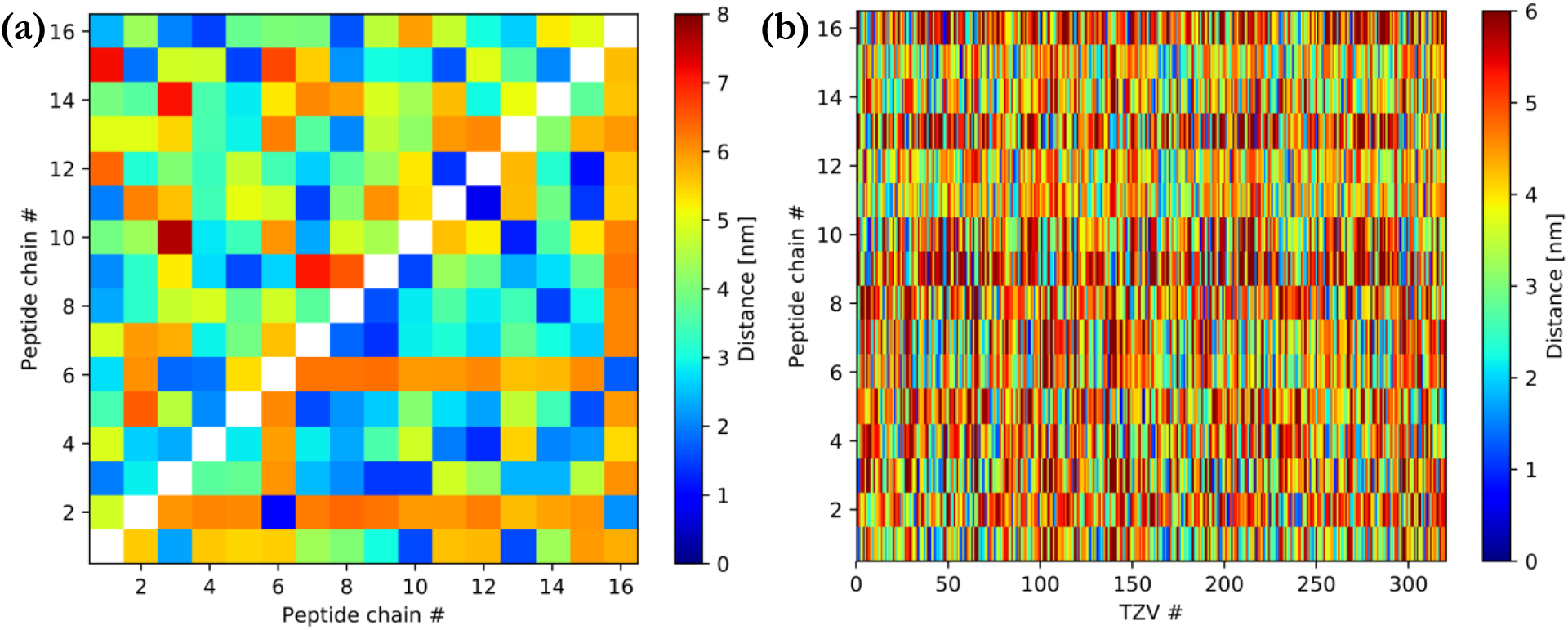
**(a)** Diagram of pairwise distances between MLT peptide molecules in the process of simulation in the presence (above diagonal) or absence (below diagonal) of 20-fold TZV excess. Without TZV, there are single pairs of molecules, sometimes close to each other (represented in dark blue). In the presence of TZV, the distance between the peptides centers of mass decreases, the number of approaching pairs increases, and clusters form. **(b)** Diagram of mean pairwise distances between MLT peptide molecules and TZV molecules during the last 100 ns of simulation. The distance is stable in time, which indicates the static position of TZV relative to MLT.

From the first moment of modeling, TZV (which has already formed chains) converges with MLT, mainly through interactions with positively charged amino acid residues (a.a.r.) and also with N-terminal a.a.r. The situation does not change substantially as the modeling proceeds (Figure 4).

**Figure 4.**
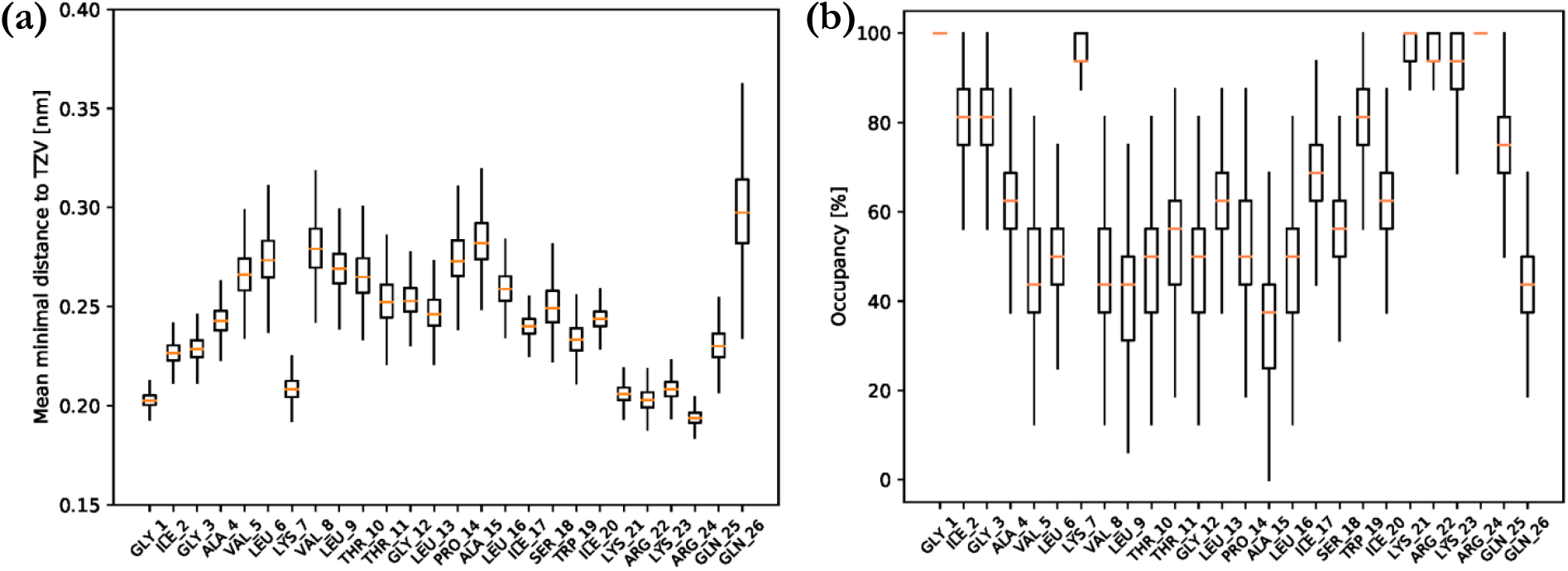
**(a)** The average minimal distances between MLT a.a.r. and TZV. **(b)** The percentage a.a.r., belonging to 16 MLT molecules, occupied by TZV molecules (the distance is lower than 0.25 nm).

It should be noted that in our earlier work [6], the antiaggregation effect of TZV chains on SI peptide (GDIRIDIRIDIRG) is also due to the interaction of TZV with positively charged a.a.r. The distances from the a.a.r. were calculated for this situation to the nearest TZV molecule (Figure 5a). The occupation of SI a.a.r. by TZV molecules were also calculated (Figure 5b), and the results are similar to those seen with the MLT-TZV system. This indicates that the mechanisms of interaction of the TZV chains and peptides are similar and are due to binding to positively charged a.a.r including N-terminal a.a.r. This supports the hypothesis that we have suggested previously [5].

**Figure 5.**
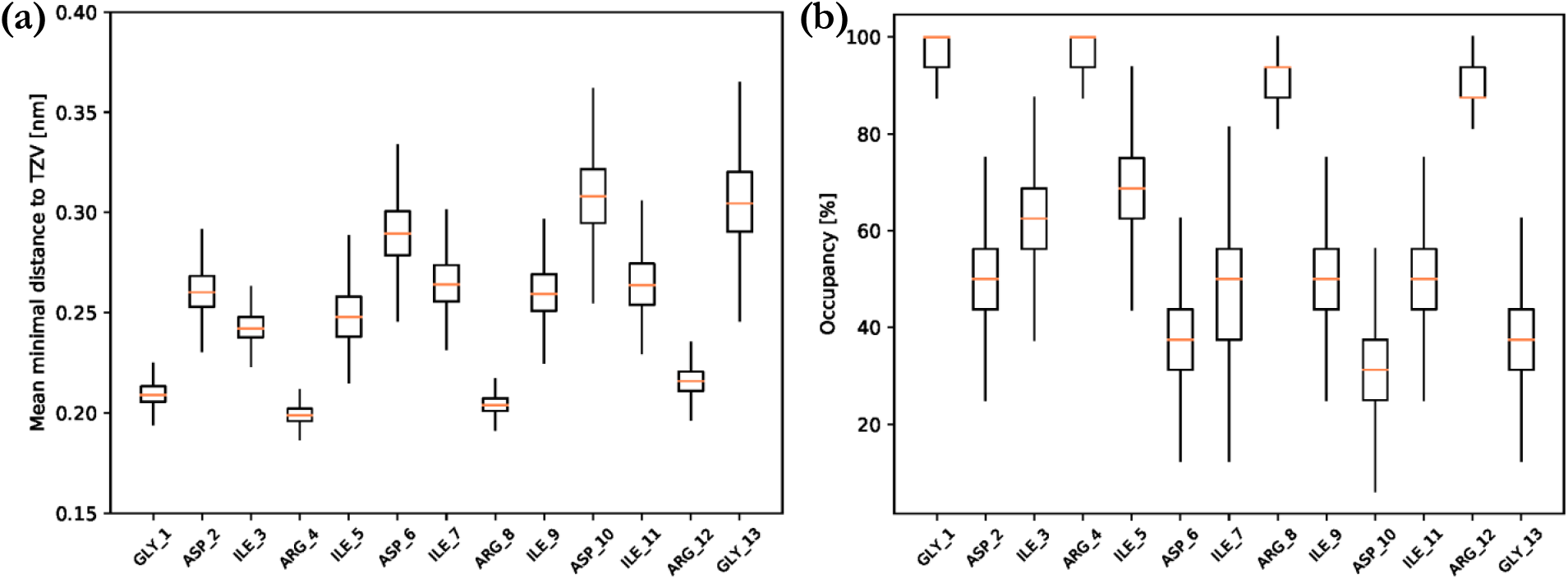
**(a)** The average minimal distances between SI a.a.r. and TZV. **(b)** The percentage of SI a.a.r, occupied by TZV molecules (the distance is lower than 0.25 nm).

According to MD data, MLT is located in a grid of TZV chains. At the same time, the system is in dynamic equilibrium. Therefore, some molecules from the surface of the aggregate can be exchanged and exhibit activity in solution. Due to this fact, precipitated MLT can retain its hemolytic activity after adding red blood cells, which is precisely what we see in the experiment.

It should be noted that during the formation of the precipitate, a small quantity of TZV was captured by MLT. It was shown by optical density measurement of the supernatant at a wavelength of 315 nm (the maximum of TZV absorption) before and after the addition of MLT. In samples with MLT: TZV ratios of 1:25, about 6 TZV molecules are recruited by one MLT molecule. Thus, induction of MLT aggregation occurs under the influence of supramolecular complexes of TZV.

A docking procedure was used in order to clarify the mechanism of interaction of TZV with a single MLT molecule (to determine the most probable binding sites with the MLT surface, and to estimate the interaction energy). Analysis of the results showed (Figure 6) that when binding to the MLT surface, TZV is most likely localized in the region containing the WIKRKR motif. In this case, TZV is located on MLT’s surface and several interactions occur. A nitro-group of TZV forms ionic and strong hydrogen bonds with Arg22. The aromatic part of Trp19 is located in the immediate vicinity of the aromatic part of TZV; this likely leads to a stacking interaction between them. The S-CH_3_ tail of TZV tends to contact with nearby isoleucines lying in neighboring helices.

**Figure 6.**
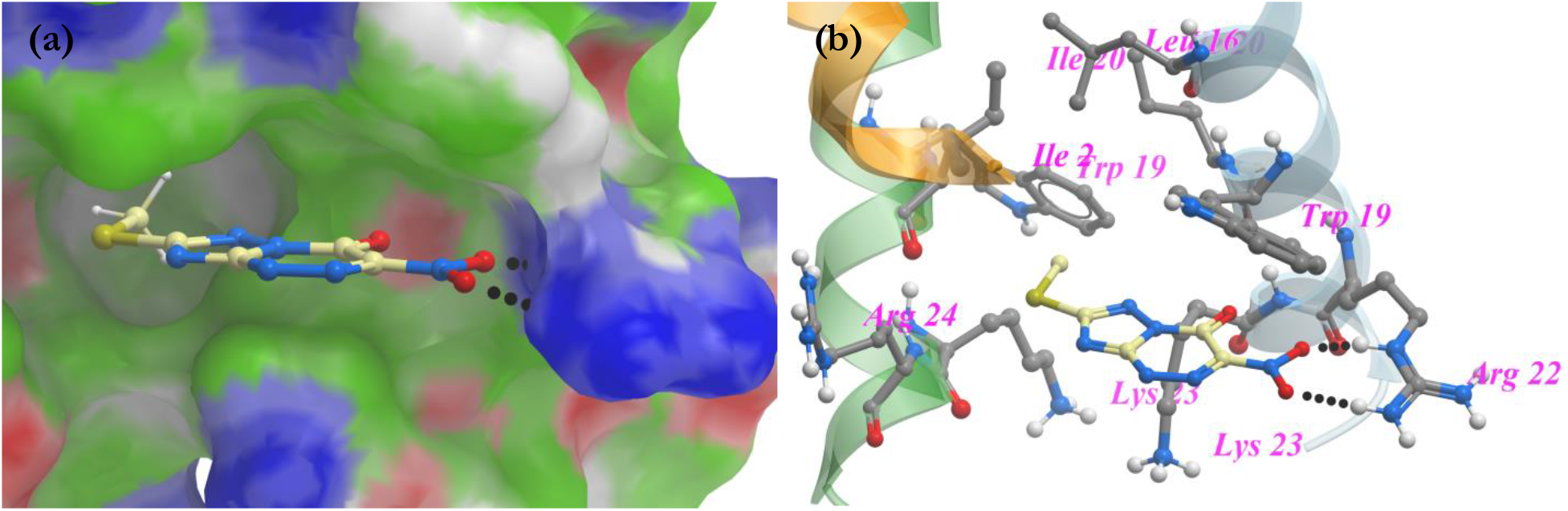
Positions of the best energy conformations obtained in the docking procedure. **(a)** TZV is located on the surface of MLT. The coloring of the surface is as follows: green – hydrophobic zones, red – hydrogen bond acceptors, blue – hydrogen bond donors, white – the surface formed by carbon atoms. Hydrogen bonds are given by dashed lines. **(b)** A.a.r. located at 5 Å from TZV. Only polar hydrogens are shown.

The energy estimations gave the following results (Table 1). The main contribution to the binding energy is provided by the electrostatic contribution. The Van der Waals energy for a given geometry of TZV on the MLT surface is not so significant. This is due to the fact that the plane in which the aromatic component of TZV is located at an angle to the plane in which Trp19 lies. The π-π interaction between them is not particularly strong. The contribution of the entropy component is insignificant because of the small conformational capacity of TZV.

**Table 1.** Values of contributions to binding energy for the complex with the best binding energy obtained by docking. All values are estimated in kcal/mol.

| $\Delta G_{\text{bind}}$ | $\Delta G_{\text{eq}}$ | $\Delta G_{\text{solv}}$ | $\Delta G_{\text{VdW}}$ | $\Delta G_{\text{hbond}}$ | $-T\Delta S$ |
| --- | --- | --- | --- | --- | --- |
| -40.11 | -44.05 | 19.84 | -16.70 | -2.35 | 3.04 |

The details of the interactions mentioned (TZV with positively charged amino acid residues) explain the tendency of MLT to aggregate in the presence of TZV. The interaction also fits with the mechanistic explanation of the antiamyloidogenic effect of TZV in arginine shielding, which we have described [5].

The combination of molecular dynamic modeling and molecular docking performed allows us to hypothesize that TZV monomers in the supramolecular complex permit binding to arginines, while the presence of TZV monomer chains provides multicenter binding. Such interactions can be more effective than binding of TZV monomers to peptide monomers [28]. As analysis of the trajectories shows, the formed supramolecular complexes interact with the MLT and SI peptide monomers near the cationic a.a.r. side chains, yet retain TZV chain structure. Weak interactions between the monomers of TZV ensure the formation of a linear structure. Such a chain, interacting even with a single amino group of the peptide, has an increased probability of finding another amino group near the first. Thus, the probability of interacting with the amino group of the peptide in the TZV molecule is higher in the formed supramolecular complex than in the monomer. Such a mechanism of supramolecular complex multicenter binding can be characteristic not only of TZV, but also of other chemical compounds, in particular, peptides.

## Conclusions

Supramolecular complexes of TZV are able to influence the oligomerization of peptides containing positively charged amino acid residues. That is to say, they promote MLT peptide aggregation and SI peptide fibril dissociation, apparently due to charge shielding. The formation of supramolecular complexes allows TZV to perform multicenter binding to cationic a.a.r. in peptide monomers.

## Acknowledgment

The results of the work were obtained using computational resources of Department of Information and Computational Technologies of the St. Petersburg State Polytechnic University (http://www.spbstu.ru). We thank Mr. Edward Ramsay for help in writing the manuscript and Dr. D.V. Lebedev (DMRB PNPI NRC KI) for fruitful discussion of the experiments.

